# SimBSI: An open-source Simulink library for developing closed-loop brain signal interfaces in animals and humans

**DOI:** 10.1101/769679

**Authors:** Alejandro Ojeda, Nathalie Buscher, Pragathi Balasubramani, Vojislav Maric, Dhakshin Ramanathan, Jyoti Mishra

## Abstract

**Objective:** A promising application of BCI technology is in the development of personalized therapies that can target neural circuits linked to mental or physical disabilities. Typical BCIs, however, offer limited value due to simplistic designs and poor understanding of the conditions being treated. Building BCIs on more solid grounds may require the characterization of the brain dynamics supporting cognition and behavior at multiple scales, from single-cell and local field potential (LFP) recordings in animals to non-invasive electroencephalography (EEG) in humans. Despite recent efforts, a unifying software framework to support closed-loop studies in both animals and humans, is still lacking. The objective of this paper is to develop such a neurotechnology software framework.

**Approach:** Here we develop the Simulink for Brain Signal Interfaces library (SimBSI). Simulink is a mature graphical programming environment within MATLAB that has gained traction for processing electrophysiological data. SimBSI adds to this ecosystem: 1) advanced human EEG source imaging, 2) cross-species multimodal data acquisition based on the Lab Streaming Layer library, and 3) a graphical experimental design platform.

**Main results:** We used several examples to demonstrate the capabilities of the library, ranging from simple signal processing, to online EEG source imaging, cognitive task design, and closed-loop neuromodulation. We further demonstrate the simplicity of developing a sophisticated experimental environment for rodents within this environment.

**Significance:** With the SimBSI library we hope to aid BCI practitioners of dissimilar backgrounds in the development of, much needed, single and cross-species closed-loop neuroscientific experiments. These experiments may provide the necessary mechanistic data for BCIs to become effective therapeutic tools.

## 1. Introduction

A brain-computer interface (BCI) is usually defined as a bidirectional communication channel between a brain and a computer to compensate for dysfunctional neuronal activity [1], rehabilitate motor skills [2], or enhance a cognitive ability [3]. Likewise, a closed-loop BCI can be defined as a BCI that uses stimulation and feedback loops to adapt its internal parameters to ongoing changes in brain dynamics, task goals, and environmental signals. A promising application of closed-loop BCI neurotechnology to clinical neuroscience is the development of personalized computerized therapies that can target specific neural circuits linked to mental or physical disabilities [4, 5, 6]. Most common BCI applications, however, offer limited therapeutic value due to oversimplified designs and poor understanding of the neurobiological basis of the conditions being treated [7, 8].

Building BCI therapies on more solid grounds may require the characterization of the brain dynamics supporting cognition and behavior at multiple scales, from single-cell and local field potential (LFP) recordings in animals to non-invasive electroencephalogram (EEG), functional near-infrared spectroscopy (fNIRS), and functional magnetic resonance imaging (fMRI) in humans [9]. For instance, taking advantage of complementary species-specific methods, cross-species studies have shed light on the neurophysiological mechanisms of fear and anxiety [10, 11, 12], rapid learning [13], and error-related adaptive learning [14]. BCI training has also shown to diminish distractibility in aging rats and humans [15].

Despite recent efforts [16], software supporting cross-species closed-loop studies is still lacking. In this paper, we introduce Simulink for Brain Signal Interfaces (SimBSI), an open-source library and auxiliary programs to facilitate closed-loop BCI research in rodents and humans. Simulink is a mature graphical programming environment for modeling and simulation of dynamical systems and signal processing that is tightly integrated with MATLAB (The MathWorks, Inc., Natick, Massachusetts, USA). Simulink has gained popularity in recent years for implementing EEG [17, 18, 19], and single-cell [20] BCIs. SimBSI adds to this ecosystem: 1) advanced human EEG source imaging methods for monitoring ongoing cortical activity, 2) flexible cross-species multimodal data acquisition based on the Lab Streaming Layer (LSL) library [21], 3) as well as a flexible experimental design platform for developing closed-loop BCI systems.

The rest of the paper is organized as follows. In Section 2 we explain the design principles that we observed for developing the SimBSI library. In Section 3 we explain how Simulink works in the context of real-time signal processing applications. In Section 4 we provide an overview of multimodal, multirate data acquisition, and synchronization using the LSL library. Sections from 5-8 comprise a series of examples for signal processing, EEG source imaging, cross-species cognitive task design, and closed-loop neuromodulation. Finally, in Section 10 we describe a hardware/software environment for closed-loop animal BCIs.

## 2. Design principles

Over the last two decades, we have witnessed a rapid development of miniaturized, low-cost bio-sensing technologies [32] accompanied by hardware of increased computing power. These advancements have fueled the creation of several software environments dedicated to human BCI where practitioners can implement a variety of predefined as well as customized approaches [33]. Ideally, one would like to extend existing human BCI systems, such as those reviewed in Table 1, to support cross-species experiments; however, this may be a cumbersome process and in many cases unfeasible due to differences in hardware, data modalities, experimental paradigms, and programming languages.

**Table 1:** BCI environments reviewed for this paper compared against SimBSI.

| Software | Language | Graphical prog. | Built-in LSL | Android and iOS | Raspberry Pi | Arduino | Ref. |
| --- | --- | --- | --- | --- | --- | --- | --- |
| BCI2000 | C++ | ✗ | ✗ | ✗ | ✗ | ✗ | [22] |
| AsTeRICS | Java, C/C++ | ✓ | ✗ | ✗ | ✗ | ✗ | [23] |
| OpenViBE | C++ | ✓ | ✓ | ✗ | ✗ | ✗ | [24] |
| BCILAB | MATLAB | ✗ | ✓ | ✗ | ✗ | ✗ | [25] |
| Pyff | Python | ✗ | ✗ | ✗ | ✓* | ✗ | [26] |
| TOBI | C++ | ✗ | ✗ | ✗ | ✗ | ✗ | [27] |
| BCI++ | C/C++, C#, MATLAB | ✗ | ✗ | ✗ | ✗ | ✗ | [28] |
| xBCI | C++ | ✓ | ✗ | ✗ | ✗ | ✗ | [29] |
| BF++ | C++ | ✗ | ✗ | ✗ | ✗ | ✗ | [30] |
| gumpy | Python | ✗ | ✓ | ✗ | ✓* | ✗ | [31] |
| SimBSI | MATLAB, C/C++ | ✓ | ✓ | ✓ | ✓ | ✓ | This paper |

\*Although it is not specified by the authors, a simplified version of this software may run on a Raspberry Pi board.

SimBSI is not a BCI software per se, i.e., it does not offer popular predefined approaches, nor is it optimized with a specific data modality in mind. Instead, we designed it as a library that implements functionality for biosignal acquisition and processing that complements the broad set of tools already available in Simulink for online signal processing (see Fig. 1). In its design we observed the following principles:

**Figure 1:**
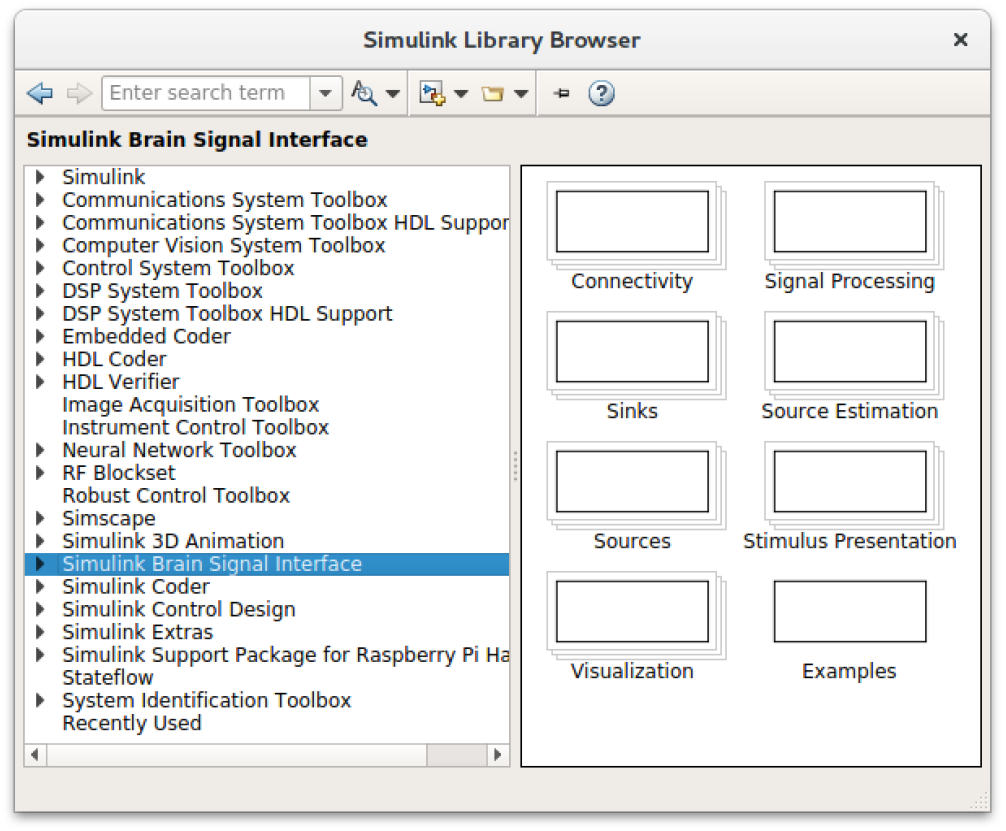
SimBSI shown inside the Simulink library browser. The documentation, code, and examples can be found online at https://bitbucket.org/neatlabs/simbsi/wiki/Home.

- Intuitive programming environment: many BCI practitioners come from fields where mastering a programming language is not a requirement. So, not having to code in the traditional way, but graphically, automatically lowers the technical background required to design a BCI.
- Transparency of data processing: the flowchart nature of the Simulink language makes it straightforward to debug and document a processing pipeline.
- Multiplatform: most Simulink programs can be compiled into standalone apps and deployed to 1) traditional platforms such as Windows, Mac and Linux, 2) embedded hardware such as Arduino and Raspberry Pi, 3) mobile devices such as Android, iPhone, and iPad.
- Flexible data acquisition: Simulink’s Instrument Control Toolbox allows interfacing instruments using standard communication protocols such as TCP/IP, UDP, and serial port communication.
- Reuse as much code as possible: large frameworks require much effort to develop, maintain, document, and validate. Since Simulink offers a well validated general-purpose online signal processing environment, why start from scratch? Furthermore, Simulink can be extended with new blocks using C/C++ or standard MATLAB code, so it is possible to reuse a large set of neuroscientific tools already developed and tested in MATLAB. For example, popular human BCI approaches could be ported from BCILAB [25] or BioSig [34] toolboxes.

## 3. Simulink in a nutshell

In this section we briefly outline how Simulink works. This is important because it will help us to understand the capabilities as well as limitations of Simulink as a platform for closed-loop BCI and how to circumvent them. We base our exposition upon what we have learned from MathWorks’ online documentation and we refer the reader to it for more in-depth explanations.

To write a program in Simulink, we use its graphical editor to connect different blocks to form a flow diagram (or pipeline) that represents the time-dependent operations performed to the signals that propagate through it. Once the pipeline is designed, we set the sample time at which the system will operate (Model Configuration Parameters *→* Solver *→* Fixed-step size); this is usually set as the inverse of the highest sampling rate of the signals acquired by the pipeline. We need to set also the stop time in the toolbar of the editor (also exposed in Model Configuration Parameters *→* Solver *→* Stop time); this is the total time that the system will run. For testing, the stop time could be any number of seconds, and for deployment, we can set it to a large number, e.g., 3600 if we want it to run for 1 hour or *inf* if we want it to run indefinitely.

In a closed-loop experiment, it is paramount to control the rate at which a pipeline is executed so that we can deliver feedback to the subject during the targeted neural state. To make sure that a pipeline runs in real-time, during the prototyping phase of a project it is a good practice to use the performance tools in the editor (Analysis *→* Performance Tools), e.g., processing 10 seconds of data should take up to 10 seconds but no more. Depending on the complexity of the pipeline it may be possible to achieve real-time performance in the default mode of operation, which is called “normal mode”. In the normal mode, the pipeline can be executed combining compiled and interpreted blocks, thus facilitating rapid prototyping, testing, and debugging. For *soft real-time^‡^* deployment, however, we would typically use an accelerated mode [35]. In accelerated modes, all blocks are translated into C code and the pipeline is compiled into a binary.

If additional performance is needed beyond what can be achieved in accelerated modes, there are two more options: the Simulink Desktop Real-Time (SDRT) Toolbox and Simulink Real-Time (SRT). SDRT technology works by turning a general-purpose computer into a real-time system during the execution of a Simulink pipeline. It does so by installing a real-time kernel that takes over the CPU at run-time so that I/O and other time-critical operations can be performed without sharing CPU time with unrelated processes. After completing each execution step, the kernel yields the CPU to serve the processes put on hold during pipeline execution. SRT, on the other hand, needs a host and a target computer. The host is used to design and tune the pipeline before it is deployed onto a dedicated (target) hardware, which can be a high-performance computer, FPGA, or DSP board for processing high volumes of data at shorter sample times. This technology also allows routing signals from the target back to the host computer for online visualization without compromising real-time performance.

## 4. Multimodal data acquisition with LSL

As we mentioned earlier, Simulink offers several blocks for standard instrument communication. For instance, in a closed-loop experiment, a pipeline could read in LFP/EEG samples via the TCP/IP protocol, filter them, extract neural markers, and send out a feedback stimulation signal if a target cognitive state is detected. If in addition, we need to save signals to disk or account for time delays between different sensors or stimulation devices, a common practice is to time stamp every sample using the clock of the LFP/EEG amplifier. To record experimental and behavioral events, it is a common practice to use TTL pulses connected to the auxiliary channels of the amplifier.

In experiments involving the collection of multiple data modalities at different rates, stimulation devices, and control signals, the synchronization via TTL pulses can become nontrivial. In these cases, a much simpler approach is to link data acquisition, stimulus presentation, and stimulation hardware with the LSL library. LSL is built on top of the TCP protocol and is designed to synchronize different devices that publish their data on the same local area network (LAN). Each sample received by the library gets assigned a time stamp calculated taking into account the offsets of the different clocks participating in the experiment. Furthermore, the time synchronization mechanism implemented in LSL was designed after the widely used Network Time Protocol (NTP) to achieve sub-millisecond accuracy. Thanks to its well-documented API, LSL has gained popularity within the EEG community and it is now supported by several hardware makers [36].

To our knowledge, so far LSL is virtually unknown in the field of animal electrophysiology. We believe, however, that its use will significantly simplify complex animal experiments. Also, recording data using the same underlying technology and file formats would enable the use of common software tools for online and offline data analyses in cross-species studies. For these reasons, we have written an LSL plugin for the Open Ephys platform [37] (see Fig. 2). Open Ephys is a popular open-source plugin-based framework mostly used for multichannel animal electrophysiology [16]. This plugin is not part of the SimBSI library but is an essential component of the LSL ecosystem that enables the development of closed-loop animal experiments using Simulink and the tools in SimBSI.

**Figure 2:**
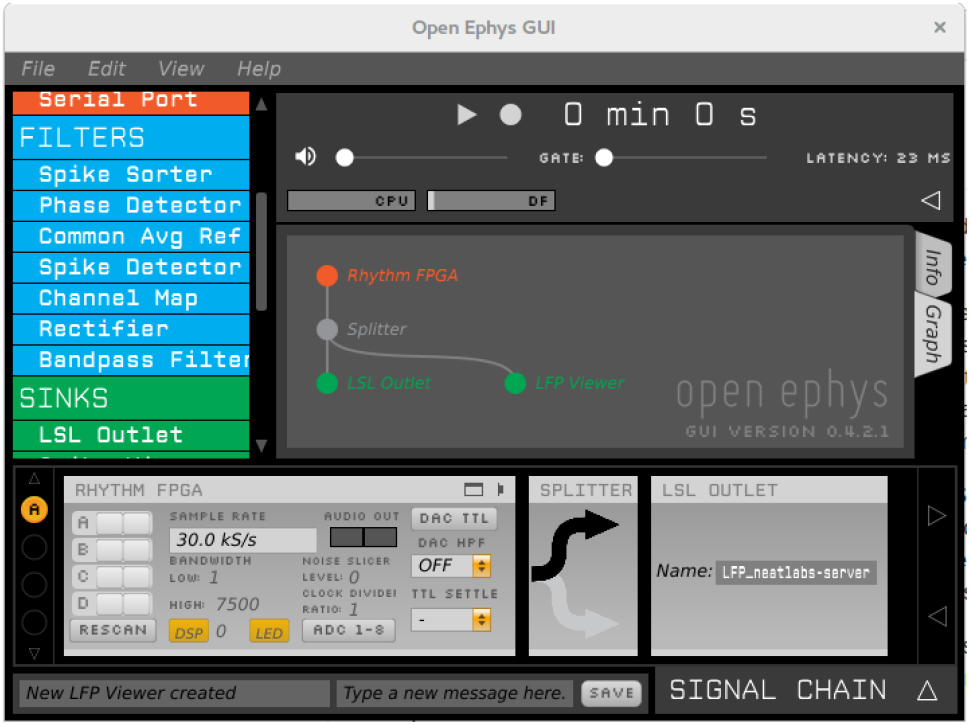
LSL plugin for Open Ephys GUI. The configuration shows Open Ephys connected to the RHD2000 FPGA Rhythm board (Intan Thechnologies, Inc., USA) for 30 kHz data acquisition from implanted electrodes and visualization. Every small batch of data acquired from the board is forwarded to LSL for online analysis or storage to disk.

To access LSL data streams within the Simulink environment, we created the *LSLInlet* block. The *LSLInlet* block is a Simulink extension function (s-function) written in C using LSL and Simulink’s APIs. Fig. 3 A shows an example of EEG data acquisition using the LSLInlet block, which can be found in SimBSI *→* Sources. To configure the LSLInlet block, we double click on it and set the stream parameters in the configuration window (panel **B**). For convenience, all the parameters can be filled up automatically by clicking on the *Select stream* button and selecting the desired stream from the list of all streams available in the LAN we are connected to (panel **C**). Panel **D** shows a 2 second snapshot of EEG data flowing through the pipeline. After the sampling rate is set, i.e., to 128 Hz, the *LSLInlet* block automatically sets the sample time of the pipeline to 0.0078125 (1/128) sec. We note that the use of the *LSLInlet* block forces Simulink to wait until a new sample is available in LSL, thereby guaranteeing an effective execution time equal or higher, but not faster than the sample time of the EEG stream.

**Figure 3:**
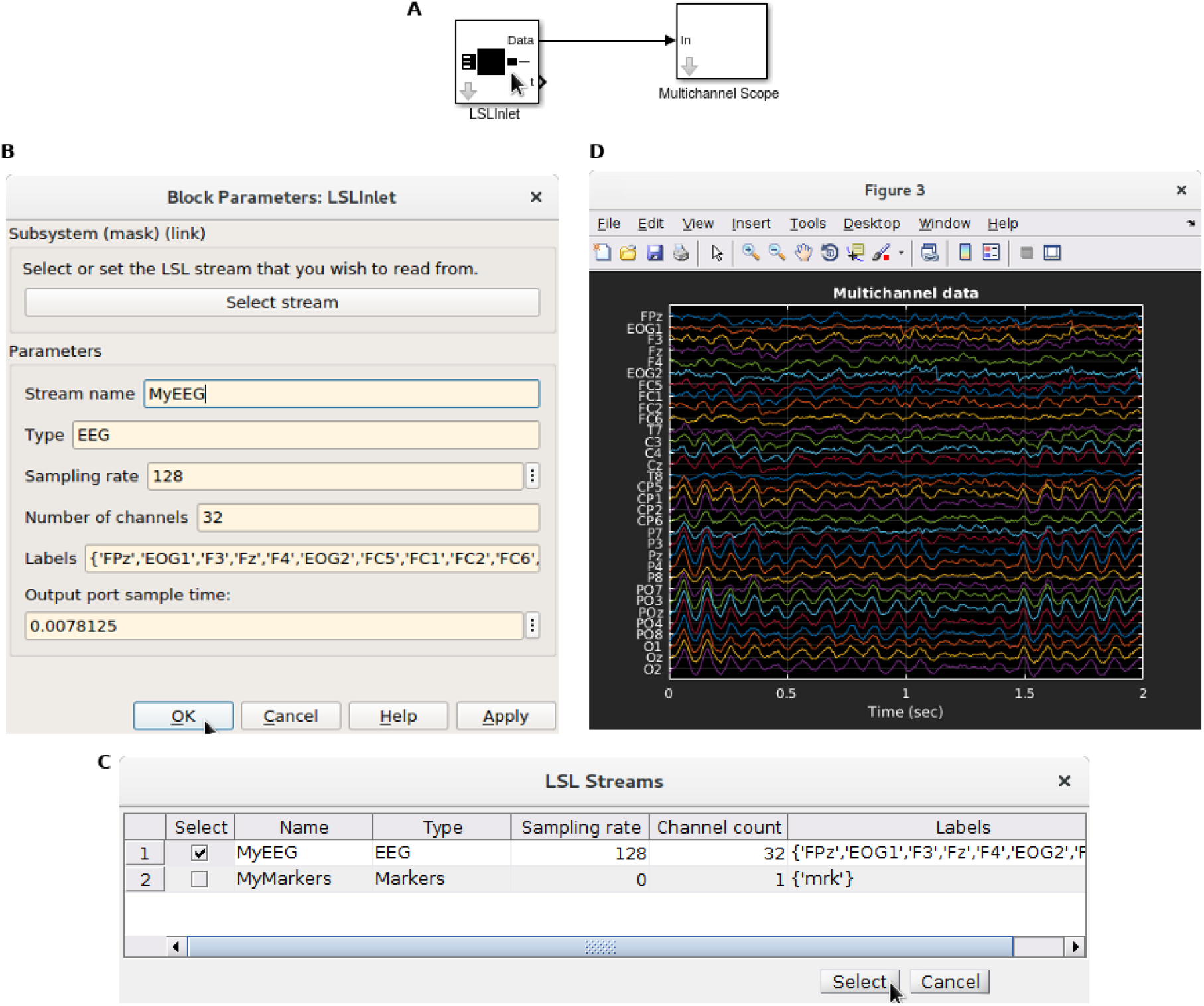
EEG data acquisition and visualization. **A**: Simulink pipeline that uses the *LSLInlet* block for reading data in from the network and the *Multichannel Scope* block for visualization. **B**: **LSL** configuration window. **C**: Stream selection tool that pops up if we click on *Select stream*.

## 5. Signal processing example

In this example, we use prerecorded EEG data to show filter design tools available in Simulink relevant to closed-loop systems. Digital filtering is a necessary step in almost all biosignal processing pipelines that requires careful consideration [38, 39]. FIR filters usually need higher orders than their IIR counterparts to satisfy their frequency response, which can incur significant time delays. An FIR filter has a linear phase response, which is desirable because, within the band of interest, the filtered signals are delayed by the same amount, thereby preventing phase distortions. Time delays can be overlooked when the analysis is performed offline. In real-time systems, however, it is worth considering the use of IIR filters at the cost of introducing small phase distortions. As we show next, the DSP System Toolbox has tools that allow us to design digital filters precisely quantifying these properties.

In Fig. 4, we used the pipeline on the left to illustrated time delays introduced by band-pass FIR and IIR filters between 1 Hz and 30 Hz. We used the block *From EEGLAB* to read prerecorded data stored in an EEGLAB-compatible file format [40] to simulate an EEG headset. This block reads all the data stored in a file at the beginning of the pipeline execution and then emits a multichannel sample on every sample time, where the latter is set automatically as the inverse of the sampling rate declared in the file. This block also guarantees that the pipeline is executed at the sampling period of the EEG by blocking the execution until that time has elapsed. Although filter blocks can work with multichannel data, in this example we use a *Selector* to process only the Fp1 channel. We use the Fp1 channel as it contains an eye-blink event to illustrate the delays introduced by each filter. Next, we filter data in parallel using FIR and IIR filters. We also compensate the raw signal by the delay introduced by each filter, concatenate raw and filtered signals and visualize them in a *Scope* block. On the right panel, we show the filter design window that pops up if we double click on a *Bandpass* block. After designing a filter, we can inspect its frequency response by clicking on the *View Filter Response* button on the top right. We show the frequency response of both FIR and IIR filters in Fig. 5.

**Figure 4:**
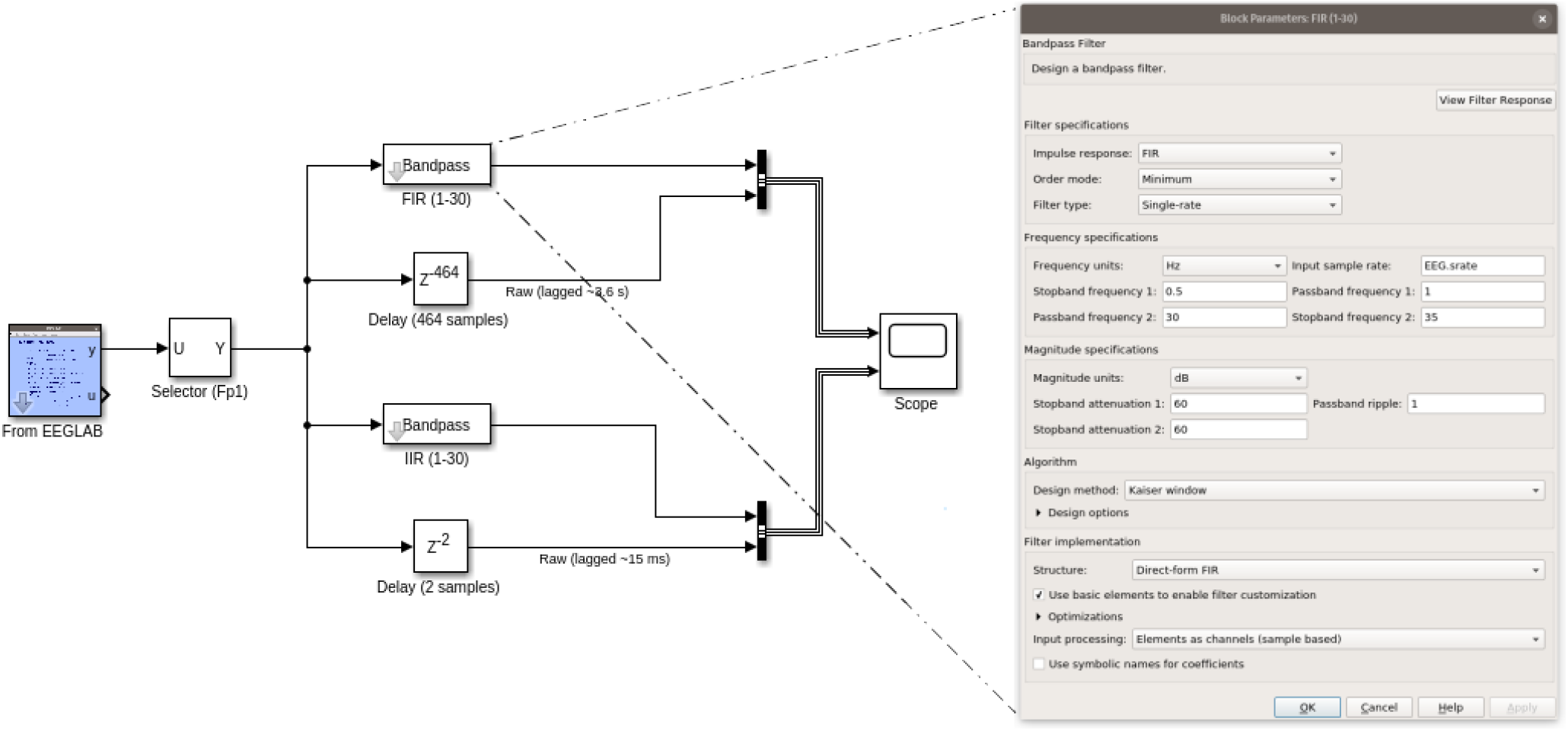
Bandbass filtering example in Simulink using blocks from SimBSI and DSP libraries. **Left**: Processing pipeline using prerecorded data. **Right:** Filter design window.

**Figure 5:**
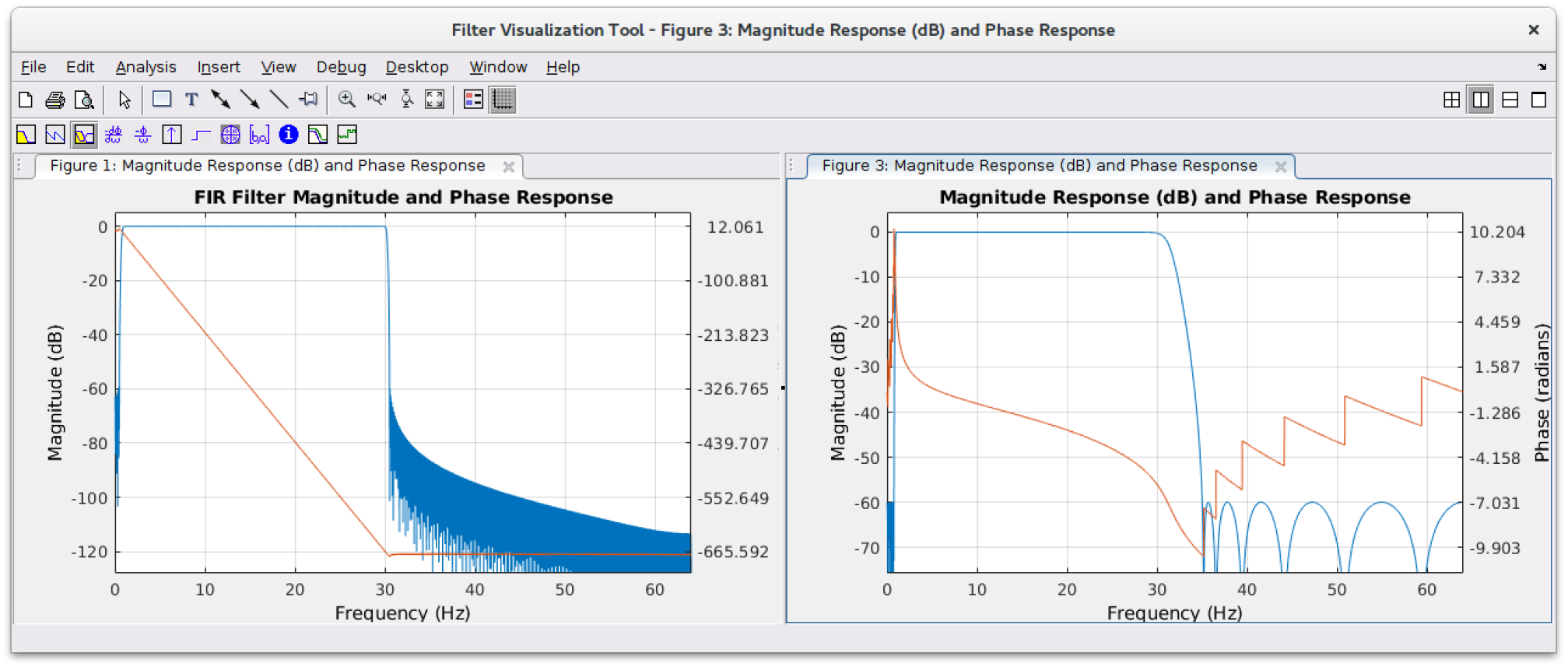
Frequency response of FIR and IIR filters used in the example of Fig. 4.

Fig. 6 shows a 10 sec window of the pipeline output in a *Scope* window. Both panels show the filtered and raw signals in yellow and blue respectively. The top panel shows the FIR branch of the pipeline and the bottom panel shows the IIR one. Note that we shifted raw signals in time by the group delay of each filter so that we could visually inspect the distortions introduced. We can see that both filters smooth the data and kept the eye-blink event. Although the FIR filter did not distort the signal significantly, it introduced a delay of *∼* 3.6 sec. Such long delay is prohibitive in a closed-loop system that reacts to brain or behavioral responses in the order of hundreds of milliseconds. The IIR filter exhibited a shorter delay of *∼* 31 msec, although we can see some distortion around the high amplitude eye-blink artifact, which may be a reasonable solution for some applications. We note that the frequency content of eye-blink artifacts overlaps with neural signals, so they typically cannot be removed by digital filters alone, thereby calling for more sophisticated artifact removal algorithms [41, 42].

**Figure 6:**
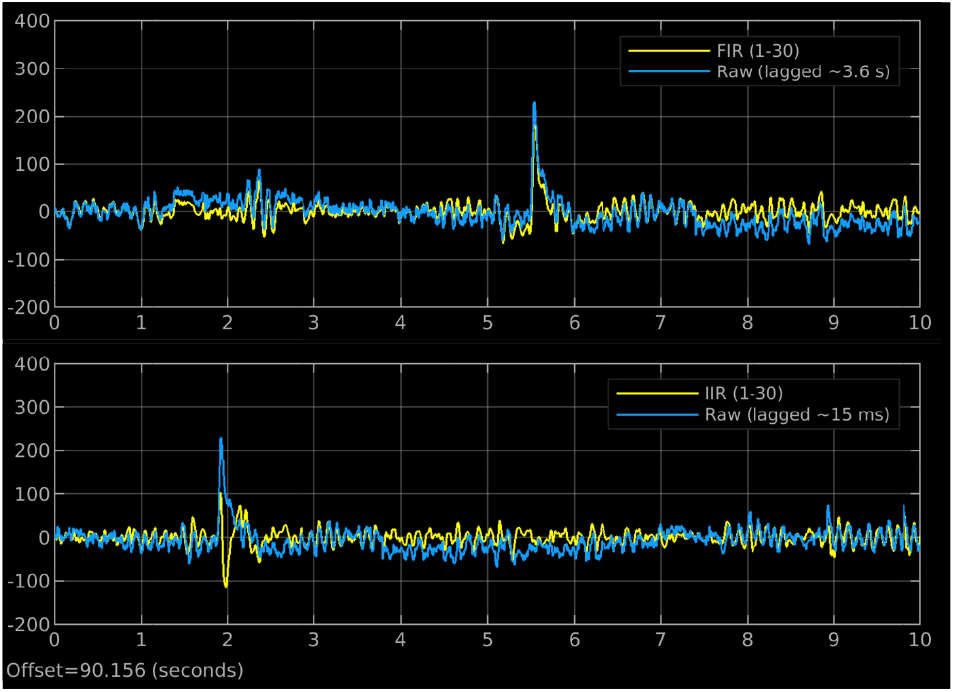
10 sec snapshot of the output of Fig. 4 pipeline. The top and bottom panels show the effect of FIR and IIR filters respectively. The eye-blink event was used to illustrate the time delay introduced by each filter.

## 6. Online EEG source imaging

It is well known that the inverse mapping from EEG scalp sensors to cortical currents is not unique. Thus, there are many instances where it is more desirable to build a human closed-loop around signals coming from specific brain regions rather than scalp sensors. For example, several studies have pointed out the therapeutic effect of transcranial magnetic stimulation (TMS) to the dorsolateral prefrontal cortex (DLPFC) for treating depression [43], chronic pain [44], and cocaine addiction [45], to mention a few. Most TMS protocols, however, are applied in open-loop fashion ignoring ongoing fluctuations in the functional state of the targeted brain region and this can hinder the efficacy of the treatment [46]. In this example, we demonstrate an EEG source imaging pipeline for monitoring the activity of several cortical areas in real-time. Then, in Section 9 we show a TMS example that can be used for building an EEG source-based closed-loop system.

The successful application of EEG-based imaging algorithms usually depends on the deployment of computationally expensive data cleaning preprocessing algorithms that typically run offline [47]. Recently, we proposed a stpatiotemporal filtering algorithm that allows for the use of continuous EEG signals to adaptively segregate brain source electrical activity into maximally independent components with known anatomical support, while minimally overlapping artifactual activity [41]. We call this algorithm Recursive Sparse Bayesian Learning (RSBL).

In Fig. 7, we show an EEG pipeline where the preprocessing is limited to the common average reference and a 1 Hz high-pass filter. Then, we use the *RSBL* block to minimize the effect of EOG, EMG, and single-channel spikes artifacts while simultaneously estimating EEG source activity in 8003 locations regularly scattered across the cortical surface of the brain. The *RSBL* block relies on the prior deployment of the *Co-register* block to co-register the sensor locations of the incoming EEG data (in this example assumed in the 10/20 system) with a template head model based on the Colin27 head. At initialization time, this block finds the common set of channels between our montage and that of the template, then it selects the corresponding columns of a precomputed lead field matrix*§*, which is used down the line by the *RSBL* block for brain mapping. At run-time, the *RSBL* block takes as input a multichannel sample of EEG data and produces the following quantities: **G**: 8003 cortical source amplitude vector, **V**: a vector containing the activity of several EOG, EMG and single channel spike artifacts, **Y_c_**: is a sample of the multichannel cleaned EEG signal, *λ*: an estimate of the common mode sensor noise variance, *γ*: a vector estimate of the scale (variance multiplier) of different groups of brain and artifact sources, and logE: the log evidence of the generative model being optimized, which is a quantity that can be used to measure the quality of the brain mapping procedure at any given time.

**Figure 7:**
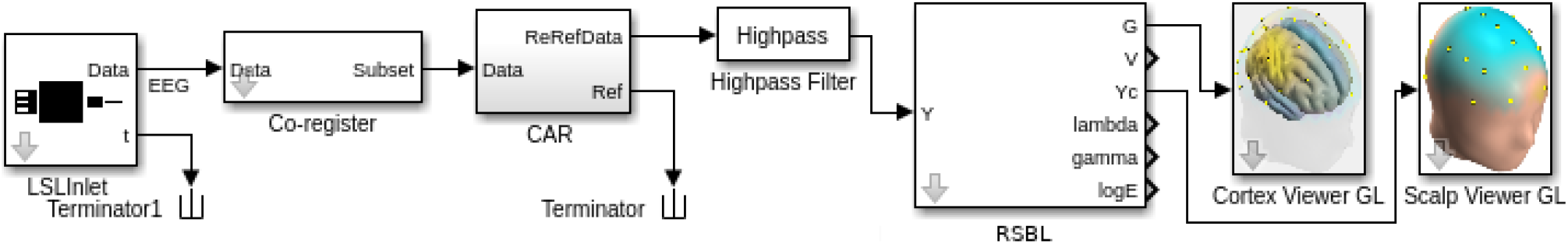
Pipeline for real-time EEG data cleaning, source separation, and imaging using the *RSBL* algorithm and OpenGL viewers.

## 7. A note about visualizations

In our experience, MATLAB-based 3D visualizations can severely hinder the real-time performance of a Simulink pipeline. So in this example, to visualize the cleaned EEG and source estimates produced by the *RSBL* block we use two freely available OpenGL-based apps written in Python, the ScalpViewer [49] and the CortexViewer [50]. The blocks *Cortex Viewer GL* and *Scalp Viewer GL* forward the data they receive to LSL which are then rendered externally by the Python apps. In the specific case of the *Scalp Viewer GL* block, it also linearly extrapolates the voltages from the sensor locations to the rest of the scalp so that we can render EEG topographies as a continuous field. We note that, in addition to LSL, the Python apps can consume data using the TCP/IP protocol, so in applications where the data need to be visualized remotely, we can simply replace the *GL* blocks in the pipeline by standard *TCP/IP Send* blocks from the Instrument Control Toolbox.

**Figure 8:**
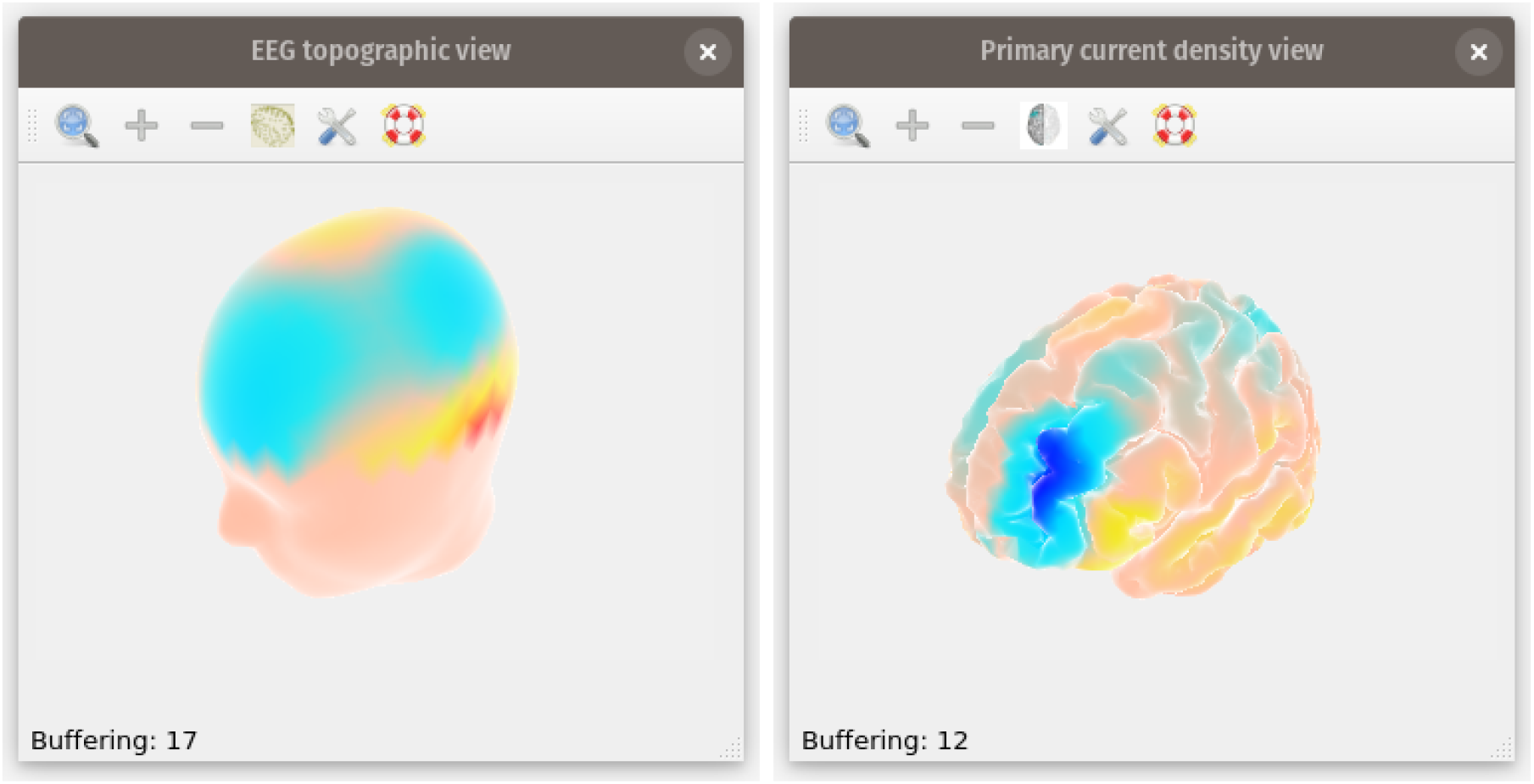
Python-based OpenGL real-time scalp and source viewers. **Left:** Scalp viewer [49]. **Right**: Cortex viewer [50].

## 8. Stateflow cognitive task design

In addition to having a fully vetted graphical environment for signal processing, of which we have full control, ideally we would like to have an experimental environment that exhibits these same features. In this section, we demonstrate the use of Stateflow for the design of cognitive tasks within the Simulink environment. Performing the experimental design within Simulink is advantageous for cross-species studies because we can implement the core logic of the task once, and then use it with species-specific data acquisition and stimulus presentation devices. Stateflow is a visual environment where we can design sequential decision logic triggered by the subject’s behavioral responses based on state machines and flow charts [51]. Furthermore, Stateflow allows us to visualize state transitions while the subject is executing the task, which is especially useful for debugging during the earlier stages of the experimental design.

In this example, we use a special type of Stateflow block called *Chart*. A *Chart* is a state machine with transitions that can be triggered by internal or external signals. Inside a *Chart*, blocks represent states while arrows represent state transitions. Each state has a name (mandatory) followed by any number of instructions written in MATLAB language; this is what the state does and is called “state action”. State actions can be of three types depending on when we want the code to be executed: 1) entry - right when the state becomes active, 2) during - during the time that the state is active, and 3) exit - right before transitioning out of the state. Here we only use entry actions because there is no obvious reason to do otherwise, but we encourage the reader to check Stateflow’s documentation for other options and use cases. A state can have any number of transitions which are then triggered in order of priority by combining logical and temporal statements. For example, in Fig. 9 B, the fourth transition of the state *WaitForResponse* is triggered after 1.8 sec of being in that state given that a “Go” image was displayed.

**Figure 9:**
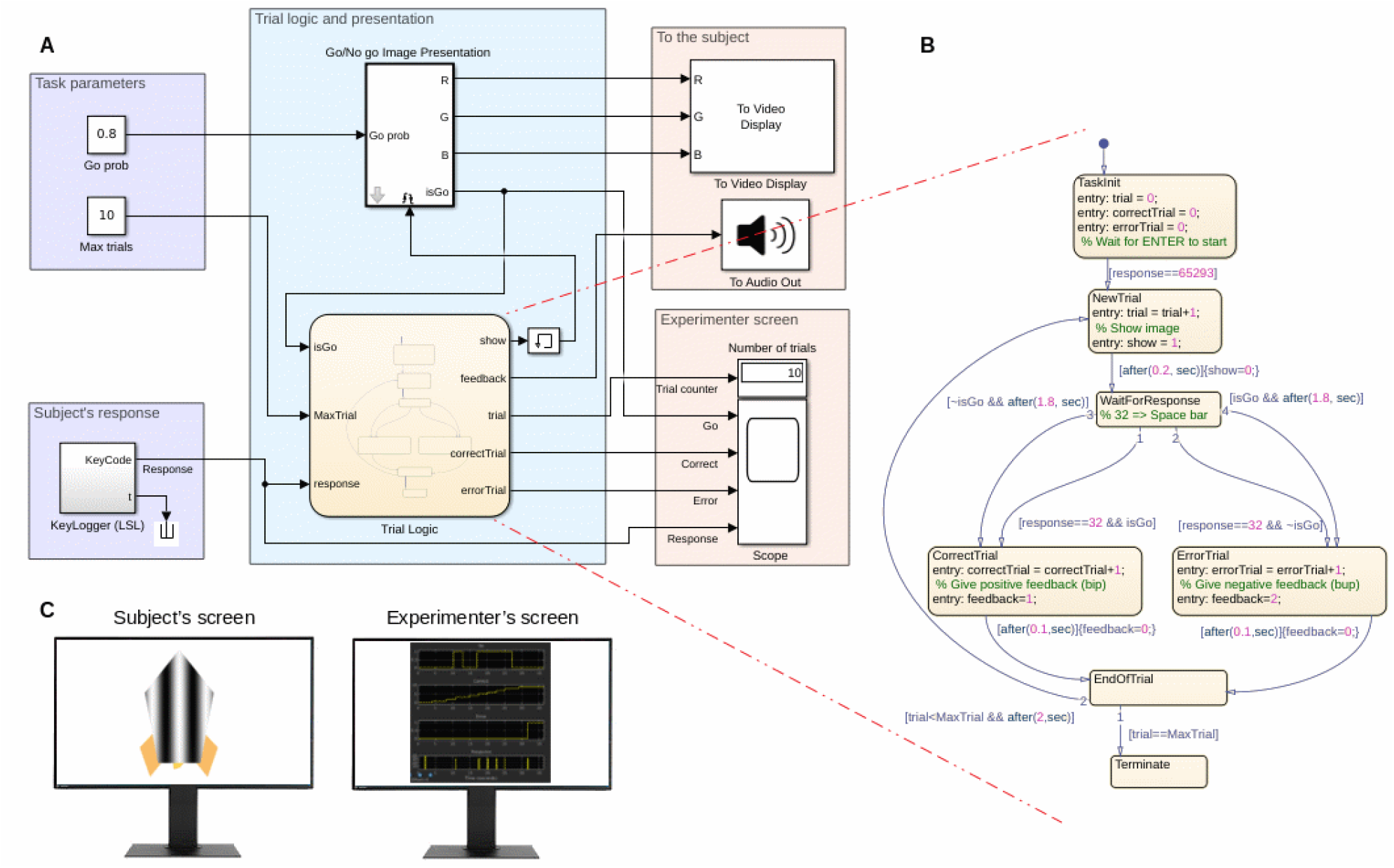
Stateflow experimental design. **A**: Parameters of the task (top) and subject’s behavior coming through LSL (bottom). **B**: Trial logic coded in Stateflow. **C**: Multimonitor setup, one for stimulus presentation facing the subject and the other one facing the experimenter in a control room.

Fig. 9 shows an example of a Go/NoGo task designed in Stateflow. Panel **A** shows the Simulink system. The blocks enclosed in the purple areas represent task parameters (e.g., Go stimulus probability and the number of trials) and the acquisition of the subject’s behavioral responses. Human subject keypresses are captured by a key logger Python app running in the background that forwards the key codes to LSL, which are then read in Simulink using the *Key Logger* block (which internally uses the *LSL Inlet* block). The blue area groups species-independent blocks for the probabilistic presentation of Go/NoGo images and the trial logic. The orange area groups visual and auditory stimuli presentation blocks as well as task and performance measures (e.g., trial counter, correct and incorrect trials, subject’s keypresses, and so on) for monitoring the progress of the task.

Panel **B** shows the *Trial Logic* block implemented as in the *Chart* block. From top to bottom, the entry point to the system is the *TaskInit* state, this is where we initialize variables and prepare for subsequent actions. To transition out of the entry point to the *NewTrial* state, and thereby start the task, we wait for the user to press the *Enter* key (code 65293). In *NewTrial* we increment the trial counter and set *show*=1; the rise of the latter triggers the *Go/NoGo Image Presentation* block outside the Chart to randomly select the image that will be immediately shown on the screen. After 200 msec, we transition to the *WaitForResponse* state and set *show*=0, which has the effect of removing the image and displaying a blank screen. *WaitForResponse* has four possible transitions: 1) to *CorrectTrial* if *response* is equal to 32 (space bar key code) and was a Go image (correct response), 2) to *ErrorTrial* if the response is 32 but was a NoGo image (incorrect response), 3) to *CorrectTrial* if there was no response for 1.8 sec and was a NoGo image (correct withholding) and 4) to *ErrorTrial* if there was no response for 1.8 sec and it was a Go image (incorrect withholding). In the *CorrectTrial* and *ErrorTrial* states we activate the respective correct or error auditory feedback tone for 100 msec and then we transition to *EndOfTrial*. We terminate after reaching the maximum number of trials. Otherwise, we wait for 2 sec and start a new trial.

Panel **C** represents a dual monitor setup in which the left screen can be used for stimulus presentation while the right one could face the experimenter in a control room. Both screens can be controlled by Simulink and all the relevant signals flowing through the system can be saved in a Matlab file using the *To File* block or sent to LSL through *LSLOutlet* blocks and then saved in the *.xdf* file format using the *LabRecorder* app [52].

## 9. EEG-based closed-loop neurostimulation

TMS has become a popular technique for studying the relationships between brain and behavior [53] through noninvasive interventions both in healthy human subjects and patients. As such, TMS have been proposed as a valuable experimental and therapeutic tool for closed-loop neuroscience [54]. For instance, it has been shown that stimulation guided by the phase and frequency of intrinsic EEG rhythms can result in different types of long-term plasticity [55]. Using this as a motivation, in Fig. 10 we demonstrate the relative simplicity of implementing a TMS closed-loop human BCI pipeline in Simulink.

**Figure 10:**
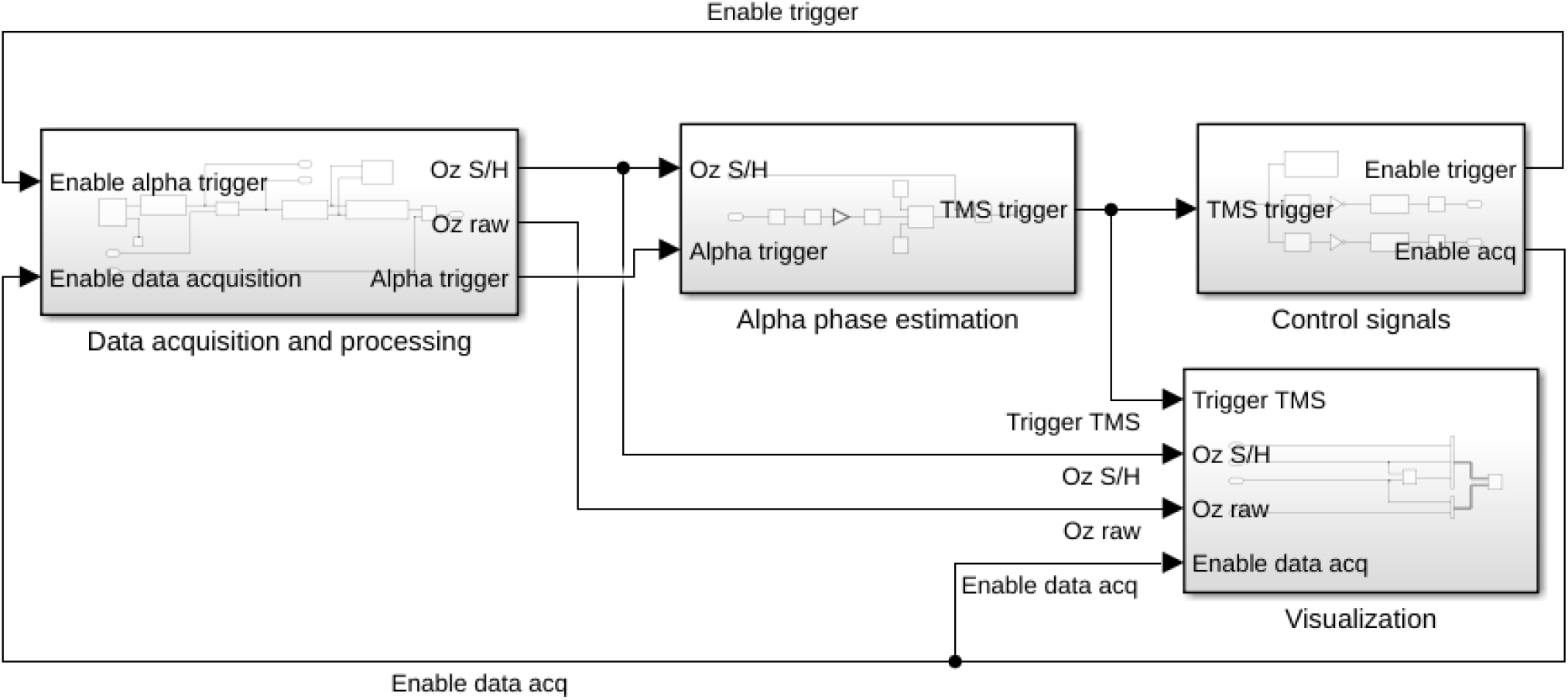
Closed-loop BCI with TMS stimulation composed of four major modules: 1) Data acquisition, 2) Alpha phase estimation, 3) Control and stimulation, and 4) Visualization. The system is designed to acquire EEG data through LSL and trigger a TMS pulse if an alpha wave is detected at the Oz channel and it is stable for at least one second. The stimulation pulse is locked to the positive peak of the EEG signal, which at that point should consist of mostly alpha rhythm activity. Each submodule is explained in the panels below.

At a higher abstraction level, the pipeline is composed of four major submodules for: 1) EEG data acquisition, 2) occipital alpha phase estimation (brain state estimation), 3) control and stimulation, and 4) visualization. The pipeline is designed to acquire EEG samples from LSL and continuously monitor the power of the alpha band at the Oz channel. When the alpha power is greater than other frequency components within 1 Hz and 40 Hz for at least 1 second, a single TMS pulse is triggered phase-locked to the positive peak of the EEG signal. Since we build on standard DSP blocks, the pipeline allows to flexibly change the channel, frequency, phase of interest, as well as the stimulation pattern sent to the TMS hardware. Figures 10-14 show in detail the different components of the system.

**Figure 11:**
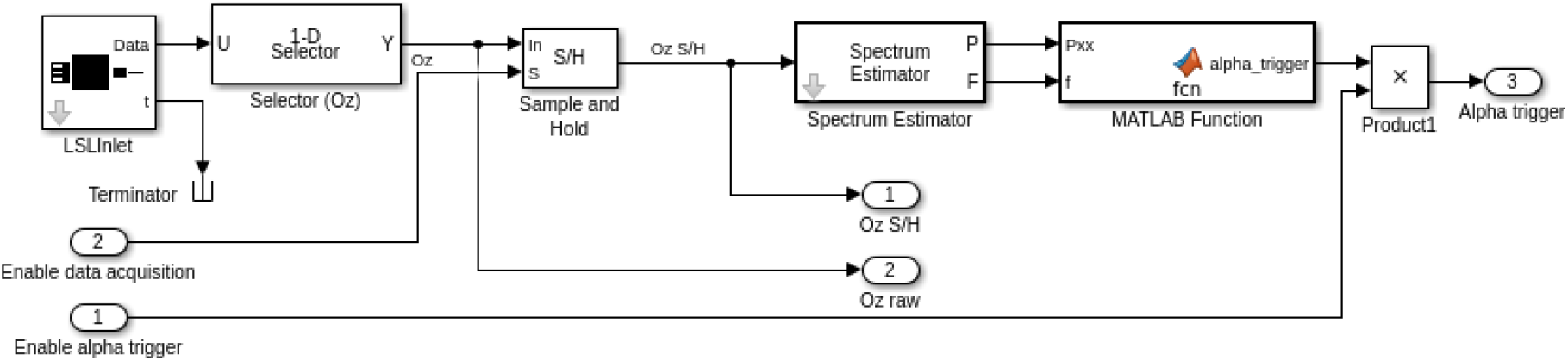
Data acquisition submodule: 1) The *LSLInlet* pulls EEG samples from LSL at the sampling rate of the EEG acquisition device. 2) The *Selector* block allows us to select the subset of channels we are going to work with, which in this case is Oz. 2) The *Sample and Hold* block enables EEG samples to pass through while the TMS pulse is not active, otherwise, it holds the last sample acquired before a TMS pulse and resumes sampling once the TMS artifacts are gone. 4) The *Spectrum Estimator* block computes the Welch (or filter bank) power spectral density estimates on 1 sec windows with a 50% overlap. 5) User-defined function that sums the power in *δ, θ, α, β, γ* bands and outputs 1 if the maximum power is in *α*, otherwise, the output is 0.

**Figure 12:**
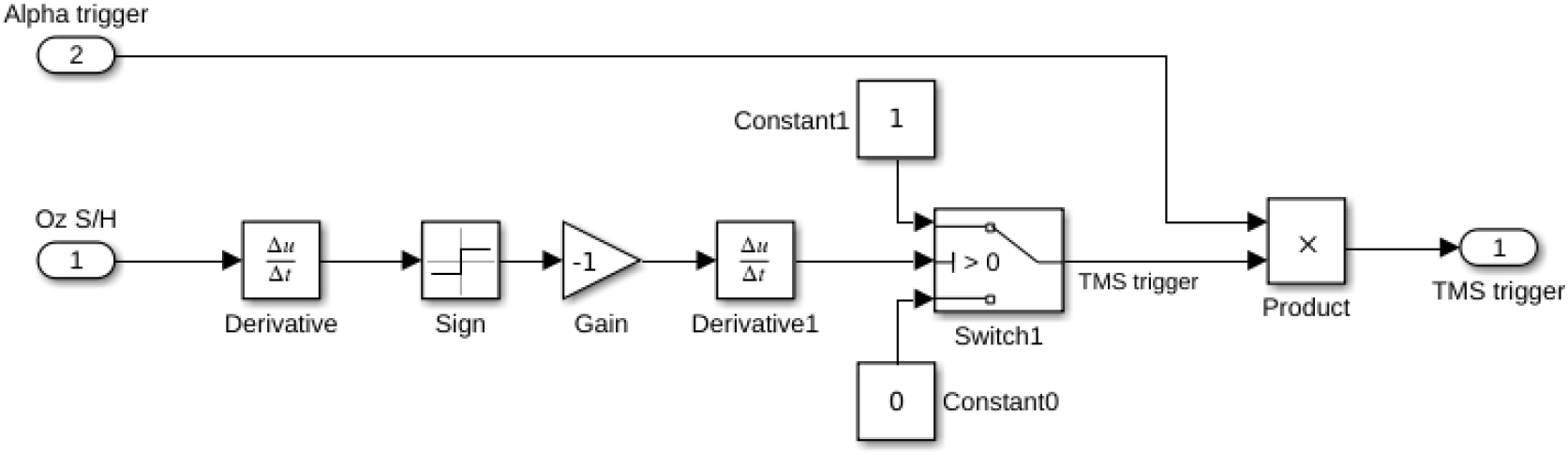
Submodule for triggering a TMS pulse time-locked to the positive phase of the *α* rhythm. The bottom branch produces a Kronecker delta at the positive peak of the Oz signal, say *y*(*t*_*k*_), implementing the expression diff(*−*sign(*y*(*t*_*k*_) *− y*(*t_k−_*_1_))). Note that to time-lock to the trough of the *α*-wave we just flip the sign of the unitary gain above.

**Figure 13:**
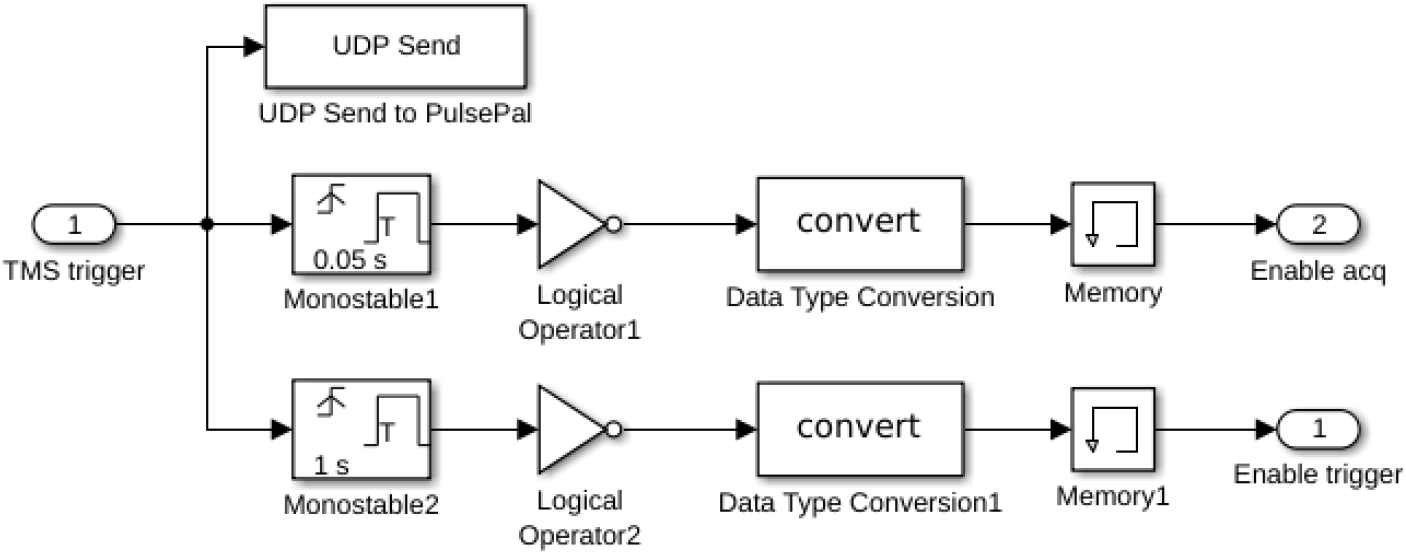
Control submodule. Once we have determined that a TMS pulse should be delivered, in parallel we send out the trigger by the serial port and use two *Monostable* blocks to 1) disable the data acquisition for the time span of the TMS artifact and 2) disable the stimulation based on *α*-waves for a prespecifed period. This can be reconfigured depending on the stimulation protocol, e.g., single pulse, burst, and so on.

**Figure 14:**
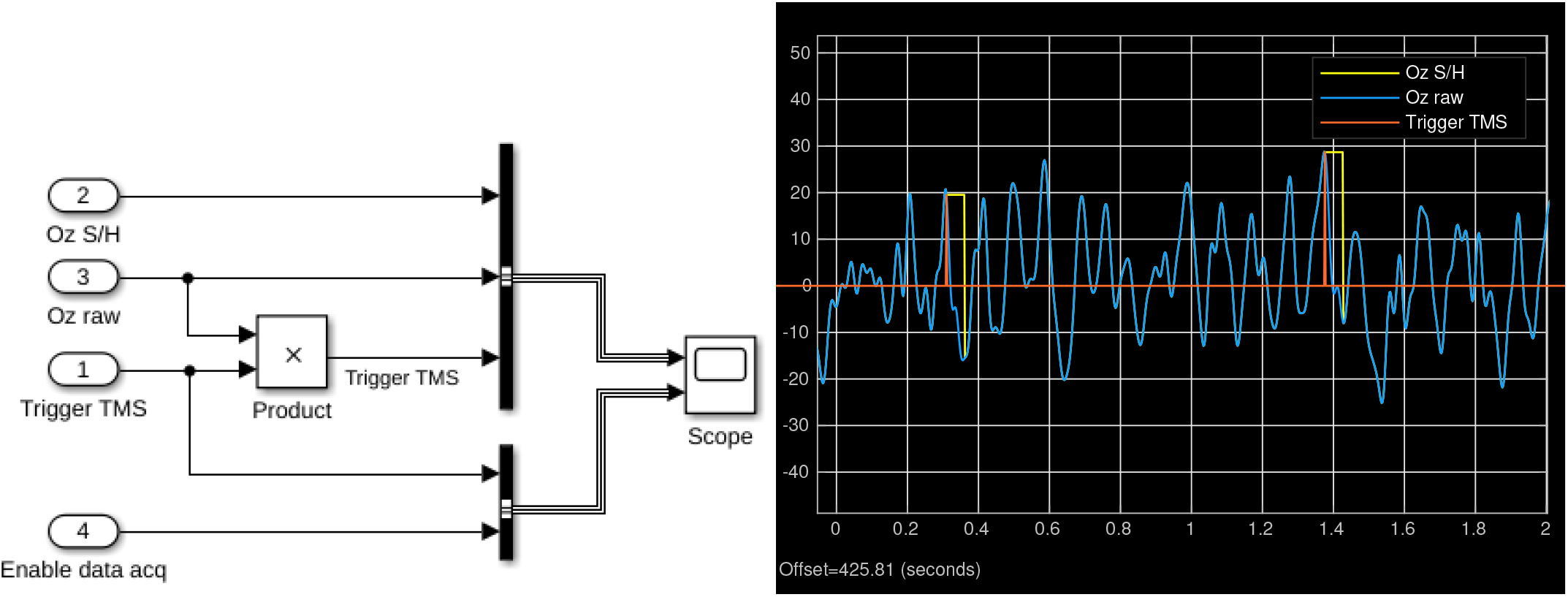
**Left:** Visualization submodule. **Right:** Excerpt of 2 sec data flowing through the system.

We send trigger pulses from Simulink to a MagVenture MagPro TMS machine using a PulsePal device [56]. PulsePal is an inexpensive and low-latency pulse generator that has been used in closed-loop electrophysiological experiments to deliver auditory and optogenetic stimulation to mice during a Go/NoGo task [57]. Furthermore, PulsePal has been recently integrated into Open Ephys. To interface PulsePal with Simulink, we used PulsePal open-source C++ API to write an s-function, which we encapsulated into the *To PulsePal* block. We note that by having a common software stack, it becomes relatively effortless to translate between human and animal closed-loop experiments, keeping most parts intact and swapping out species-specific components only, e.g., EEG by LFP signals and TMS by optogenetic stimulation hardware.

## 10. Animal experimental environment

Our last example blends most elements explained so far (with the exception of human EEG imaging) for the creation of an experimental environment for rodents. The environment consists of two major components: 1) a behavioral box and 2) an electrophysiology data acquisition system. Although this environment will be described in detail in a separate publication, next we proceed to briefly explain how we used elements in the SimBSI library and auxiliary apps to integrate Stateflow task design, behavioral and electrophysiology data acquisition, stimulation and post hoc data analysis.

### 10.1. Behavioral box

The behavioral box consists of a custom built chamber where animals are placed to perform a cognitive task while we record their behavioral responses and electropysiological signals (see Fig. 15 A). The chamber has a display where visual stimuli can be presented. In front of the display there are 5 nose ports, each of which is equipped with an infrared (IR) sensor for collecting animal choices and a liquid reward delivery mechanism. There are speakers placed on each side of the display for delivering auditory feedback and house lights glued to the top face of the chamber, which are often used to indicate to the animal the start of the task or to reinforce negative feedback in the form of short bursts of 10 Hz flashing light.

**Figure 15:**
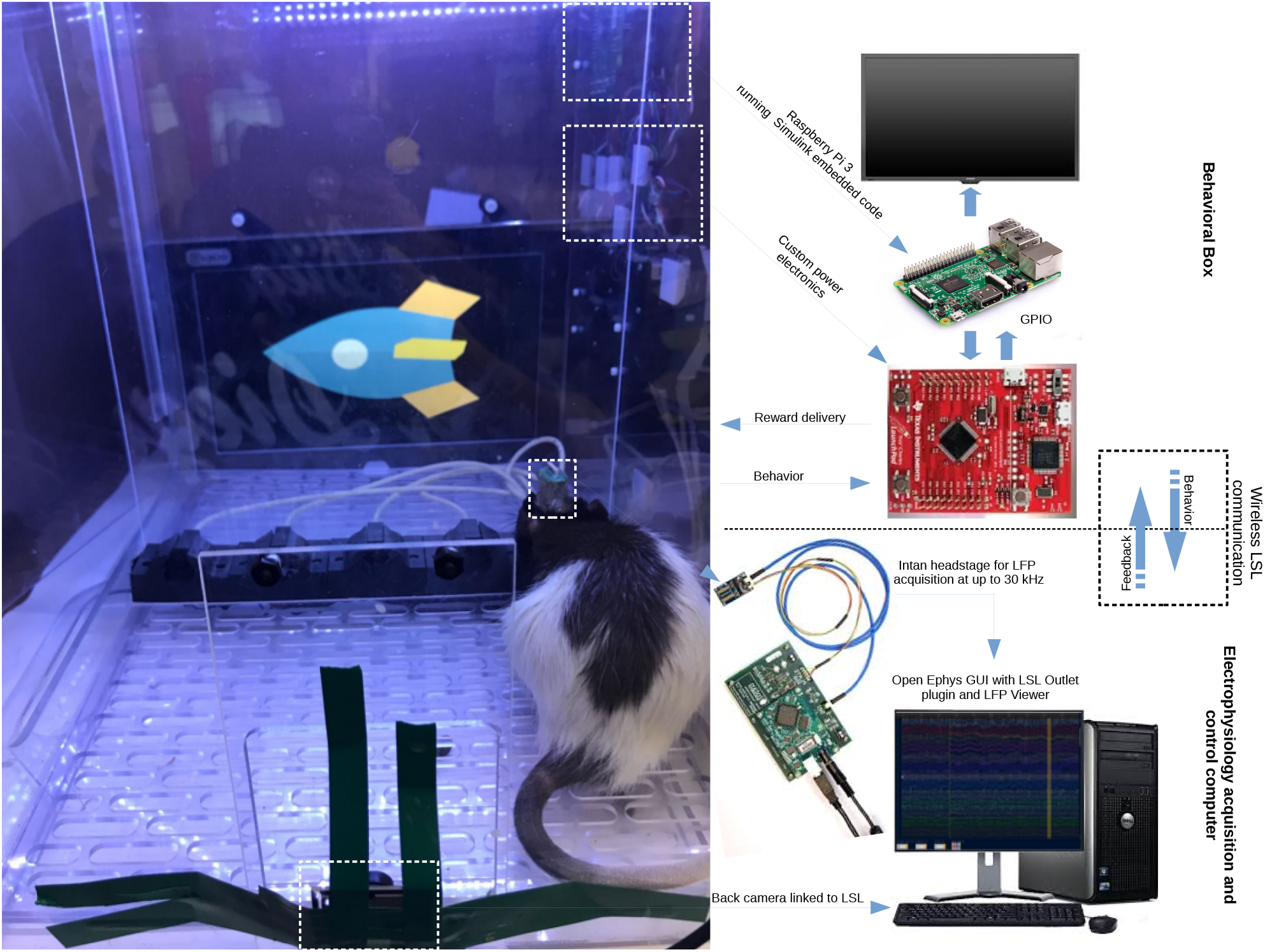
Animal experimental environment.

The display, IR sensors, reward mechanism, auditory feedback, and house light are controlled by a Simulink process running on a Raspberry Pi 3 model B+ (Pi for short) that is attached to the outer face of one of the walls of the box (see Fig. 15 B). The Pi interfaces with sensors and actuators via the GPIO pins connected to a custom-built power electronics control board. Before deployment to the behavioral box, we assign an IP address to the Pi and configure it to connect to a dedicated LAN (typically wireless), this way we can have several boxes in the lab and treat them as self-contained mobile experimental environments.

Once the Pi is fully configured and attached to the box, it can be programmed using a Simulink session running on a remote computer connected on the same network. Common cognitive tasks such as the one shown in Section 8 can be coded using Stateflow and Simulink blocks on the host computer, simulated, and then deployed to the targeted Pi using the MATLAB Coder. MATLAB Coder uses our Simulink program to generate C code that is optimized for the Pi and runs natively on it as a real-time standalone application. Once the task is being compiled, it stays permanently in the Pi’s SIM card. To access compiled tasks stored on the Pi without using Simulink, we have created a MATLAB GUI called BrainER^||^ (see Fig. 16). With BrainER we can select one or several Pis and reconfigure, stop, and start tasks on them.

**Figure 16:**
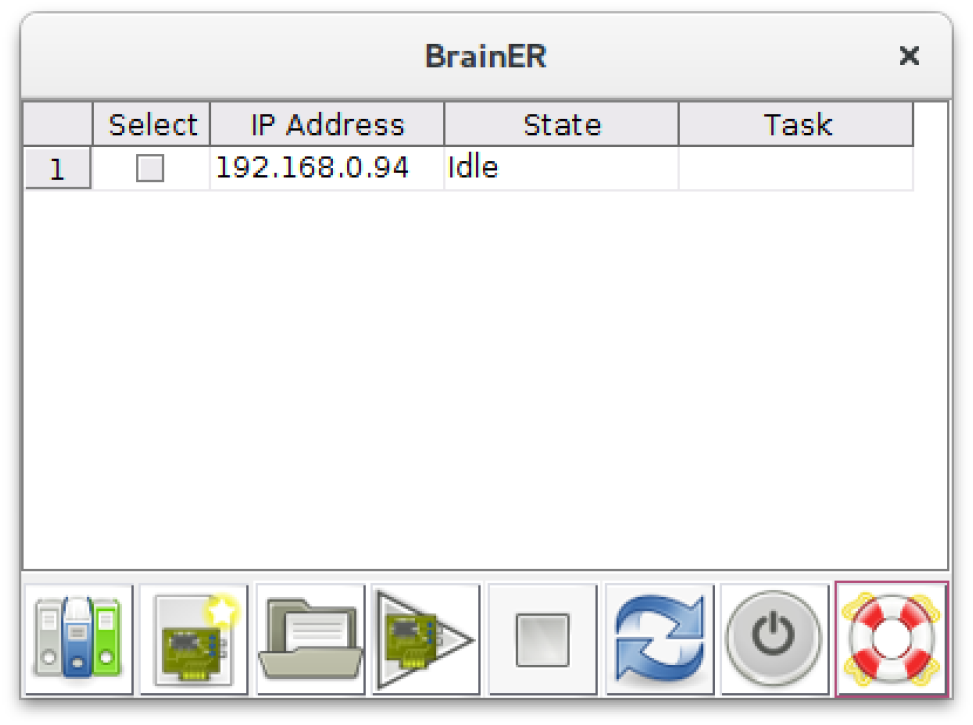
The BrainER GUI allows us to manage the software that runs on Pis that control several behavioral boxes that we may have in the lab. Once the GUI is first launched, it searches the network for Pi devices connected between IP addresses 192.186.0.90-150.

### 10.2. Electrophysiology data acquisition system

The animals that participate in the tasks administered through the behavioral box are usually implanted with intracranial electrodes. To acquire LFP data, we use Intan headstages connected to a RHD200 acquisition board, which is controlled using the Open Ephys software. To synchronize the LFP data with the control and behavioral signals coming in and out of the behavioral box we use LSL. As mentioned in Section 4, we use the *LSLOutlet* plug-in for Open Ephys to forward each LFP sample acquired to LSL. To enable LSL communication on the behavioral box, we enhance the capabilities of the read/write GPIO blocks in the Raspberry Pi support library. We use this library to create new blocks that also forward to LSL the signals they read in or write out, respectively.

Since LFP, behavioral, and control signals can be accessed via LSL, we can save the data generated on each session using the LabRecorder app, and analyze them offline in MATLAB using MoBILAB [58] and EEGLAB [40] toolboxes and custom scripts. Furthermore, in experiments where we need to close the loop, we can implement an additional Simulink pipeline that reads the LFP data from LSL, which is required to monitor the target brain state and triggers a stimulation device, similar to what we did in Section 9.

## 11. Conclusions

In this paper we have developed SimBSI, an open-source Simulink library for the rapid prototyping of brain signal interfaces. We designed the library observing the following principles: 1) easy to adopt by users with different levels of technical expertise (e.g., students, clinicians, BCI engineers), 2) transparency of data processing, 3) multiplatform, and 4) flexible data acquisition. Most of these design principles are achieved by using a mature signal processing environment such as Simulink, powered by an intuitive graphical programming language. Simulink programs are multiplatform and can be deployed as standalone applications that run on standard or embedded hardware such as a Raspberry Pi and Arduino. We extended the data acquisition capabilities of Simulink by implementing LSL streaming blocks. We used several examples to demonstrate the capabilities of the library for implementing cross-species BCIs, ranging from simple signal processing, EEG source imaging, cognitive task design, to closed-loop neuromodulation. Furthermore, we demonstrated that a sophisticated experimental environment for rodents is feasible and relatively straightforward to develop using Simulink and the SimBSI library. We hope that SimBSI will facilitate the development of, much needed, single and cross-species closed-loop neuroscientific experiments. These experiments may provide the necessary mechanistic data for BCIs to become effective therapeutic tools in the future. Users can find the documentation, code, and examples online at https://bitbucket.org/neatlabs/simbsi/wiki/Home.

## 12 Acknowledgments

This research was supported by NIMH training fellowships in Cognitive Neuroscience T32MH020002 and Biological Psychiatry and Neuroscience T32 MH18399 (AO), UC San Diego Chancellor’s Research Excellence Scholarship (JM, AO), UC San Diego School of Medicine start-up funds (JM), and the Department of Veterans Affairs, Veterans Health Administration Career Development Award 7IK2BX003308 (DR) and a Career Award for Medical Scientists from the Burroughs Wellcome Fund 1015644 (DR). The RSBL algorithm is copyrighted for commercial use (UC San Diego Copyright #SD2018-817) and free for research and educational purposes.

## Footnotes

‡ By soft real-time we mean a pipeline that is executed at a non-deterministic step size, although, it is fast enough on average to keep up with the speed of the data acquisition; this is typically the case of pipelines executed on standard multicore desktop computers. Conversely, hard real-time refers to a pipeline that runs on dedicated hardware with a strictly deterministic step size.

§ The lead field is an overcomplete dictionary of unitary source scalp projections that can be calculated by solving Maxwell’s equations in a discretized model of the head obtained from MRI data (see [48] for details).

|| https://bitbucket.org/neatlabs/brainer/wiki/Home

## Notes

https://bitbucket.org/neatlabs/simbsi/wiki/Home

https://bitbucket.org/neatlabs/brainer/wiki/Home

